# Accurate and flexible estimation of effective population size history

**DOI:** 10.1101/2024.10.16.618650

**Authors:** Zhendong Huang, Yao-ban Chan, David Balding

## Abstract

Current methods for inferring historical population sizes from DNA sequences often impose a heavy computational burden, or relieve that burden by imposing a fixed parametric form. In addition, they can be marred by sequencing errors or uncertainty about recombination rates, and the quality of inference is often poor in the recent past. We propose “InferNo” for flexible, nonparametric inference of effective population sizes. It requires modest computing resources and little prior knowledge of the recombination and mutation maps, and it is robust to sequencing error and gene conversion. We illustrate the statistical and computational advantages of InferNo over previous approaches using a range of simulation scenarios. In particular, we demonstrate the ability of InferNo to exploit biobank-scale datasets for accurate inference of rapid population size changes in the recent past. Applying InferNo to worldwide human data, we find remarkable similarities in inferences from different populations in the same region. The historical sizes of all non-African populations converge around 40 000 years ago, but they remain distinct from those of African populations for over 800 000 years, long predating estimates for the migration of modern humans out of Africa.

## Introduction

We propose a new method, “InferNo”, which takes DNA sequence data and infers effective population size history, providing clues about migrations, plagues and environmental events. Some features of InferNo:

- It is computationally and statistically efficient for the recent and distant past. In particular, it can exploit biobank-scale datasets to provide accurate inferences of recent population size changes.
- The inference is nonparametric, requiring no pre-specified population size model.
- Variation in mutation rate along the genome is accommodated, only an average needs to be pre-specified.
- An average recombination rate must be pre-specified, but InferNo is robust both to variation along the genome and misspecification of the average rate.
- It is robust to sequencing errors, requiring no information about the error rate.

Some existing approaches infer population size history from estimated times since the most recent common ancestor (TMRCA) along pairs of sequences, under the sequentially Markov coalescent (SMC) model. These include PSMC Li and Durbin (2011), MSMC Schiffels and Durbin (2014), diCal Druet et al. (2014), SMC++ Terhorst et al. (2017) and PHLASH Terhorst (2024). When the sample size *n* is small, these approaches can provide good inferences for ancient population sizes, but tend to perform poorly over recent history and are computationally demanding for large *n*. Moreover, these approaches usually require good knowledge of the mutation and recombination rate maps across the genome to accurately specify the SMC, and they can be sensitive to sequencing errors.

A second class of approaches involves using the allele (or site) frequency spectrum (AFS, or SFS) of the genome sequences Gutenkunst et al. (2009); Excoffier et al. (2013); Bhaskar et al. (2015); Kamm et al. (2020). Variation in mutation rate along the genome presents a particular difficulty for these methods, and they can also be sensitive to sequencing errors. The AFS ignores the ordering of sites along the genome and so avoids any assumption about recombination, but the resulting loss of information can lead to an ill-posed inverse problem (a given AFS can arise from different demographic histories), especially for small *n* DeWitt et al. (2021).

A third class of approaches is based on inference of the ancestral recombination graph (ARG), which summarises the genealogical history of the sequences Palamara et al. (2012); Boitard et al. (2016); Speidel et al. (2019); Fournier et al. (2023). The true ARG provides all information available from the sample about population size history. However, ARG inference is imprecise, in part due to uncertainty about recombination and mutation rates, and computational costs can be high despite recent advances Rasmussen et al. (2014); Speidel et al. (2019); Kelleher et al. (2019); Mahmoudi et al. (2022); Deng et al. (2024b).

Population-size inference can also be based on inferred identity-by-descent (IBD) relationships or linkage disequilibrium (LD) coefficients Mezzavilla and Ghirotto (2015); Browning and Browning (2015); Browning et al. (2018); Ragsdale and Gravel (2019); Santiago et al. (2020). These methods are typically limited to recent history due to the challenging nature of inferring short IBD segments and the complex relationship of LD with demography.

InferNo uses the AFS in a novel way that improves on previous approaches, and provides further improvement by adding information from pairwise TMRCA estimates. It uses a simulation under the standard coalescent model to estimate the distribution of mutation events, then identifies a time scaling of the standard coalescent to best match the observed AFS and TMRCA estimates. The inferred population size is proportional to the inverse of the time scaling.

In simulation studies, we find that InferNo is overall more accurate and computationally faster than the TMRCA-based PSMC Li and Durbin (2011) and CHIMP Upadhya and Steinrücken (2022), AFS-based Mushi DeWitt et al. (2021), ARG-based Relate Speidel et al. (2019), and IBD-based IBDNe Browning and Browning (2015). In application to human data, we demonstrate inferences of population size history that are remarkably similar for populations in the same region. We find evidence for two periods of population decline in the past 150 000 years for most non-African populations, and a single, more ancient population decline in ancestral African (AFR) populations. A substantial difference in the demographic histories of AFR and non-AFR populations dates back more than 800 000 years.

## Methods

### Overview of InferNo

The data are *n* homologous genome sequences of length *ℓ*, with alleles either 0 (ancestral) or 1 (derived); ancestral alleles can often be accurately inferred from related species. Required input also includes 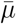, a genome-average mutation rate (per generation per site). In deriving the method we assume that, at each site, at most one mutation has occurred since the MRCA, but we demonstrate robustness to this assumption by not incorporating it into the data generation models of the simulation study.

InferNo fits a coalescent model with effective population size that is a piece-wise constant function over time. By scaling time inversely proportional to population size, this size-varying model becomes equivalent to a standard coalescent model (the “null model”) with population size constant over time. InferNo uses penalised least-squares regression to infer the time scaling and hence the population size function.

First, a simulation is performed under the null model, generating a dataset with the same dimensions as the observed data. The null-model time axis is divided into intervals, and corresponding intervals are inferred in the size-varying model by relating observed statistics to sums over time intervals of simulation-generated statistics. One statistic is the AFS, which aggregates information across sequences and across sites. The AFS can suffice for large *n*, otherwise we also use TMRCA estimates from pairs of sequences within genome segments.

Below we describe the key features of each step. More details are given in Supporting Information (SI).

### The null model and time scaling

The recombination rate *r* is assumed constant over sites and over time. The mutation rate is constant over time, but site-specific rates *µ*(*s*) can be pre-specified or estimated using a piecewise-constant smoothing of the fraction of sites that are polymorphic, scaled so that the average over all sites is 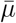 (see SI for details).

We initially assume that the observed sequences originate from a coalescent-with-recombination model with *N* (*g*), the effective population size *g* generations ago, a piece-wise constant function. This variable-size model is equivalent to the constant-size null model (with population size 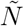) with time scaled in proportion to 1*/N* (*g*) Griffiths and Tavaré (1994). We infer the time scaling by comparing the observed dataset to a new dataset with the same sample size and sequence length generated from the null model.

First, we partition the null-model time domain into intervals 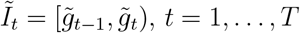, where 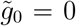. By default, we choose the 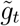 to approximately equalise the number of null-simulation mutations during each interval, but they can be pre-specified, for example, based on historical periods of interest or prior knowledge about *N* (*g*).

We then infer time intervals *I*_*t*_ = [*g*_*t*−1_, *g*_*t*_) in the variable-size model, with *g*_0_ = 0. Within each *I*_*t*_, the value of *N* (*g*) is constant and such that the cumulative coalescence rate over *I*_*t*_ is the same as for 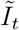. It follows that the time interval lengths are inversely proportional to population sizes, so that for *g* ∈ *I*_*t*_,

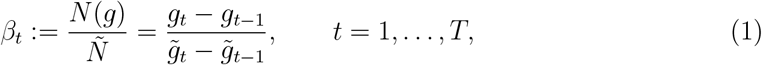

where:= indicates a definition.

The inference task is now, given our choices for 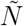 and the 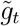, to estimate the *β*_*t*_ and hence the *g*_*t*_. The value of 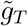 should be large enough that the inferred *g*_*T*_ predates the historic period of interest, but it should not be larger than the largest TMRCA arising in the null simulation as there is no information for inference prior to that time.

### Using the AFS

The observed-data AFS is a length-*n* vector *Y*_1_ with *k*th element equal to the number of sites at which the derived allele has frequency *k*. From the null simulation, we obtain the *n* × *T* matrix 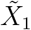 in which column *t* is an AFS restricted to sites at which the mutation arose during 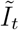 (oldest mutation if there is more than one). We can now infer *β* using the regression equation

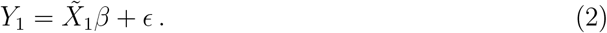

The resulting estimates are accurate when *n* is large, but for small *n* we seek additional estimating equations by relating TMRCA estimates between the observed data and null simulation.

### Using pairwise TMRCA estimates

We partition the genome [0, *ℓ*) into segments *J*_*l*_ = [*h*_*l*−1_, *h*_*l*_), *l* = 1, …, *L*, containing roughly equal numbers of variable sites. Then, for each of *n* sequence pairs we count the sites in each *J*_*l*_ at which the sequences differ in order to estimate *Y*_2_, a vector of quantiles of the pairwise TMRCA distribution averaged over segments. Next, we estimate the corresponding quantiles from our null simulation, once again separated by the time interval of the mutation (the resulting matrix 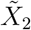 has a column for each time interval). Similar to (2), we can now infer *β* via

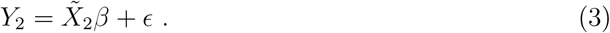

### Combining the estimating equations

We combine [2] and [3] by defining *Y* = (*Y*_1_^*′*^, *ωY*_2_^*′*^)^*′*^ and 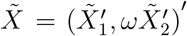, where ^*′*^ denotes matrix transpose and *ω* is a scalar weight. For large *n*, we set *ω* = 0, thus ignoring [3] which greatly reduces computational effort.

We then estimate *β* by minimising a penalised least-squares based on

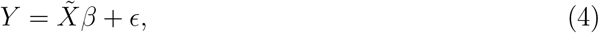

with the smoothing penalty based on the second difference of *β*, parameterised by *λ*. See SI for selection of *λ* by minimising an adjusted Bayes Information Criterion (BIC).

An optional final step is to convert the piece-wise constant estimate 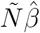 into a continuous piece-wise linear function by connecting the interval midpoints.

### Design of simulation studies

To compare the performance of InferNo with Relate, PSMC, Mushi and CHIMP, we used msprime Baumdicker et al. (2021) to generate sequences of length *ℓ* = 10^7^, with sample sizes *n* = 10 and 200, under Models 1 to 6 (Table 1). We additionally compare with PSMC for *n* = 2, corresponding to a single diploid individual. We use 25 replicate datasets for each setting, and compare performance using RMISE (root mean integrated squared error).

**Table 1:** Population sizes (in units of 1K sequences, K denotes 10^3^) at growth-rate change points for eight simulation models. Growth rates are constant between change-points, see Table 5 for the rates. A blank entry indicates no change from the value to its left. *N* (*g*) = *N* (30K) for *g >* 30K.

| Model | N(0) | N(25) | N(30) | N(300) | N(400) | N(1K) | N(1.8K) | N(1.9K) | N(3K) | N(9K) | N(18K) | N(18.1K) | N(30K) |
| --- | --- | --- | --- | --- | --- | --- | --- | --- | --- | --- | --- | --- | --- |
| 1 | 20 |  |  |  |  |  |  |  |  |  |  |  |  |
| 2 | 40 | 40 | 40 | 39 | 38 | 36 | 33 | 33 | 30 | 16 | 7 | 7 | 2 |
| 3 | 5 | 5 | 5 | 6 | 6 | 8 | 12 | 13 | 22 | 28 | 34 |  |  |
| 4 | 40 | 40 | 40 | 38 | 37 | 33 | 28 | 27 | 22 | 12 | 16 | 16 | 23 |
| 5 | 40 |  |  |  | 10 |  |  | 33 |  |  |  | 20 |  |
| 6 | 100 | 84 | 81 | 12 | 6 | 31 |  |  |  | 6 | 23 |  |  |
| 7 | 3 000 | 20 |  |  |  |  |  |  |  |  |  |  |  |
| 8 | 300 | 86 | 182 | 177 | 175 | 165 | 152 | 151 | 135 | 74 | 30 | 30 | 9 |

Models 1 to 3 have constant, monotonic increasing and monotonic decreasing population sizes at varying rates. Models 4 to 6 have periods of both growth and decline, with population size changing smoothly in Model 4 and with jumps in Model 5, while Model 6 allows us to examine inference of an older bottleneck when its signal may be blurred by a more recent bottleneck.

See SI for implementation details of the comparison methods. For the InferNo analyses, genome-wide average recombination and mutation rates were set to the correct values for the mutation and recombination maps used in the simulations (Fig. 1), but the variation along the genome is not input. To further challenge inferences, we include sequencing errors in the data simulations, with rate *ϵ* = 1 per 10^5^ sequence sites, and gene conversions with rate 2 per site per 10^8^ generations and tract length 300 sites, which are close to recent estimates for humans Williams et al. (2015).

**Figure 1.**
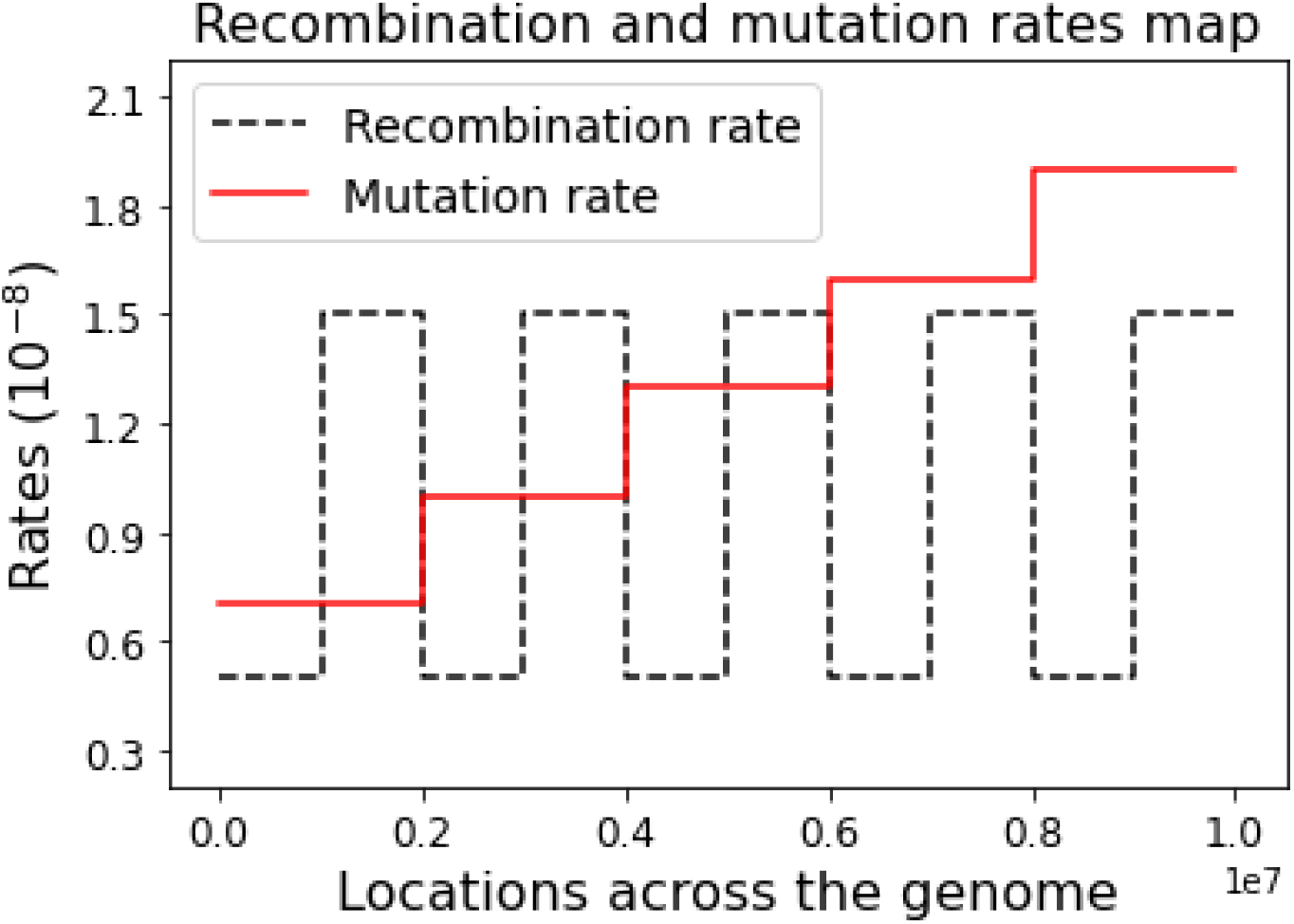
Recombination and mutation maps used in the simulation study. InferNo requires genome-wide average values as input, which for these maps are *r* = 1 and *µ* = 1.3 per site per 10^8^ generations.

Using the same 6 models, we then assessed the performance of InferNo over sample sizes (*n* = 10, 50, 100, 500, and 1000 sequences) and under two variations to the data generating model that we now describe.

In the case *n* = 100, we checked the robustness of InferNo to misspecification of the recombination rate across the genome, by simulating datasets with (i) the recombination map given in Fig. 1, which has average rate *r* = 10^−8^ per site and per generation, (ii) a constant rate *r* = 10^−8^ (which matches the InferNo analysis model) and (iii) a constant rate *r* = 1.5×10^−8^.

Again with *n* = 100, we checked the robustness of InferNo to sequencing error. We ran InferNo on simulated datasets contaminated by sequencing error with rates *ϵ* = 0, 1, 2, and 3 errors per 10^5^ sites.

To focus on recent population size inference, we compared InferNo with the IBD-based approach IBDNe Browning and Browning (2015) using Model 7 (rapid, recent growth) with no sequencing error or gene conversion. For both methods, we used *n* = 2 000, length *ℓ* = 10^8^, input the correct mutation and recombination rates *µ* = 1.3*r* = 1.3×10^−8^ and set *ω* = 0. Because of our focus on the recent past, we set time intervals for InferNo as 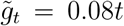 for *t* = 1, 2, 3, and 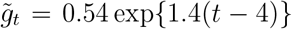 for *t* = 4, …, 12. For IBDNe, we input IBD segments of length *>* 2 cM identified by hap-ibd Zhou et al. (2020) with the genetic map parameter set to true. Default values were used for other IBDNe and hap-ibd parameters.

Finally, we tested the ability of InferNo to infer very recent population size changes (Model 8) using a biobank-scale dataset (*n* = 2×10^5^) for which IBDNe is not tractable. We set *ℓ* = 5×10^7^, similar to human chromosome 21, and kept *µ* = 1.3*r* = 1.3×10^−8^. Sequencing error is included, at rate 1 per 10^5^ sites, but not gene conversion. We set 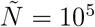 in InferNo, and to prioritize very recent inferences, 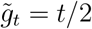 for *t* = 1, …, 50, and 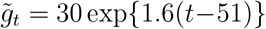 for *t* = 51, …, 56.

### Human data analysis

We estimated *N* (*g*) from all 22 autosomes for 16 contemporary human populations, including four populations from each of Africa, East Asia, Europe, and South Asia (Table 2). The samples came predominantly from the 1000 Genomes Project Fairley et al. (2020), supplemented by the Human Genome Diversity Project Bergstrom et al. (2020) and the Simons Genome Diversity Project Mallick et al. (2016). Rather than using the original sequence data, we extracted AFS and pairwise differences required for the InferNo analyses from a genealogy constructed using genome sequences from 3 601 modern and eight high-coverage ancient humans Wohns et al. (2022). Use of this unified genealogy helps ensure uniform data quality thanks to data consistency checks based on the full dataset. Our analysis used a subset of the genealogy corresponding to 1 625 modern humans.

**Table 2:** Sample sizes (haploid) for human data analysis. The number of individuals is *n/*2.

| Pop | Description | Sample size $n$ | Region |
| --- | --- | --- | --- |
| LWK | Luhya in Webuye, Kenya | 206 | Africa (AFR) |
| MSL | Mende in Sierra Leone | 180 |  |
| ESN | Esan in Nigeria | 200 |  |
| YRI | Yoruba in Ibadan, Nigeria | 214 |  |
| CHB | Han Chinese in Beijing, China | 212 | East Asia (EAS) |
| JPT | Japanese in Tokyo, Japan | 210 |  |
| CHS | Southern Han Chinese | 210 |  |
| CDX | Chinese Dai in Xishuangbanna, China | 200 |  |
| FIN | Finnish in Finland | 210 | Europe (EUR) |
| GBR | British in England and Scotland | 200 |  |
| IBS | Iberian Population in Spain | 214 |  |
| TSI | Toscani in Italia | 222 |  |
| BEB | Bengali from Bangladesh | 172 | South Asia (SAS) |
| ITU | Indian Telugu from the UK | 204 |  |
| PJL | Punjabi from Lahore, Pakistan | 192 |  |
| STU | Sri Lankan Tamil from the UK | 204 |  |
| Total |  | 3 250 |  |

For the InferNo analyses, to reduce computational effort we optimized the smoothing penalty *λ* using the chromosome 20 data only, and applied the resulting value to the analyses of all 22 autosomes. See SI for details.

## Results

### Simulation study results

For *n* = 200, Fig. 2 shows that all five methods perform relatively poorly at recent times (*g <* 10^3^) and when *g >* 3×10^4^. In the former case, the small number of recent recombination events limits the number of distinct lineages that can contribute to inference, while in the latter case, the number of lineages has been reduced by coalescences. Rapid changes of *N* (*g*) with *g* also challenge every method.

**Figure 2.**
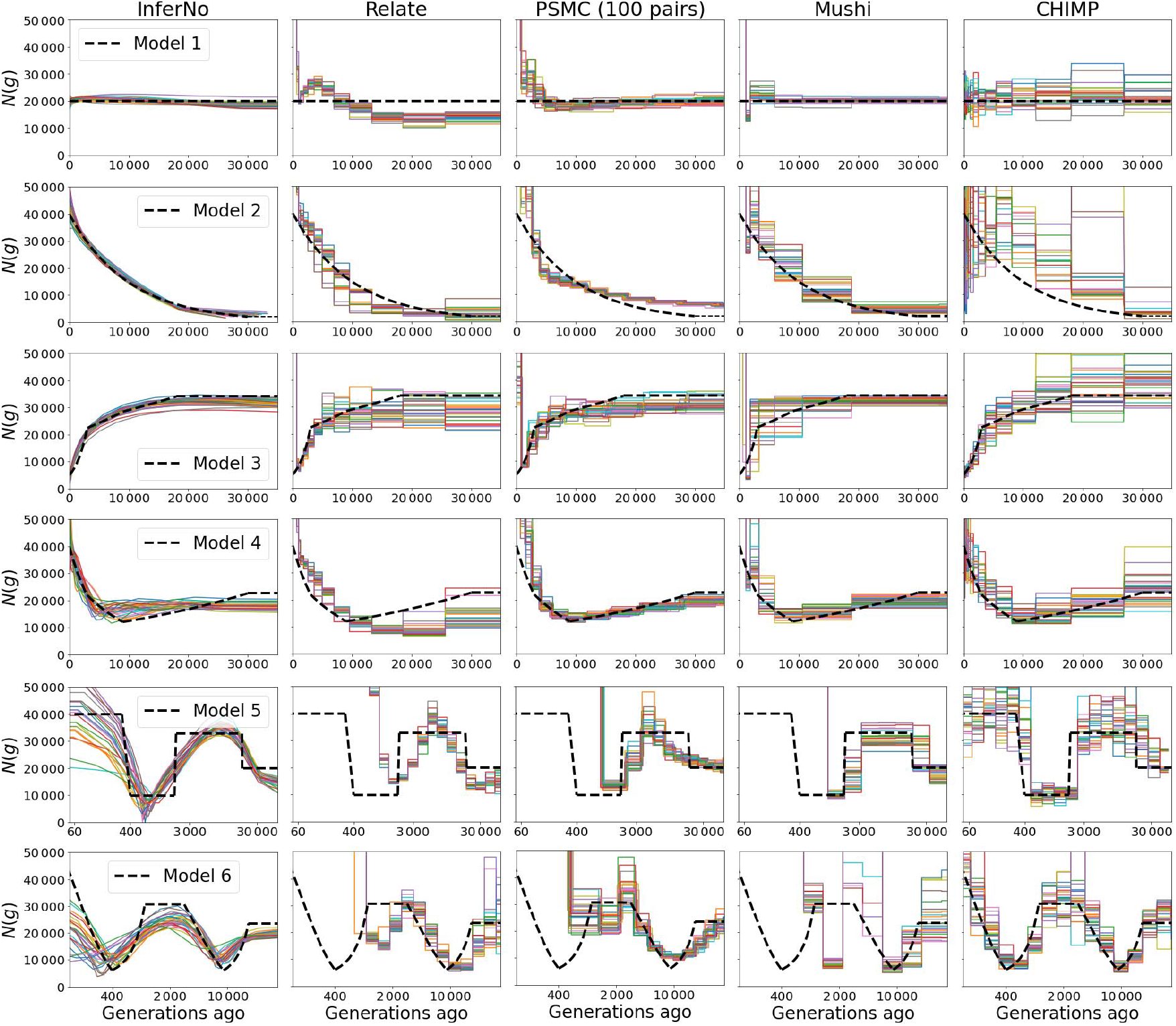
Estimates of population size *N* (*g*) in Models 1 to 6 using five inference methods. Sample size *n* = 200 and sequence length *ℓ* = 10^7^ sites. Note logarithmic time scale for Models 5 and 6. See Table 3 for corresponding RMISE values, Fig. 8 for plots when *n* = 10, and Table 6 for computing times.

On the criterion of minimising RMISE over *g* ∈ [200, 30 000], InferNo is superior to all other methods, in most cases by orders of magnitude (Table 3). This primarily reflects its superior performance for *g* ∈ [200, 1 000). When we remove this time interval and re-compute RMISE for *g* ∈ [1 000, 30 000], the methods become more comparable, but InferNo is still the best for all six models. In the special case of a single diploid individual (*n* = 2), InferNo greatly outperforms PSMC for *g <* 5 000 (Table 4), while for *g >* 5 000 the two methods show similar accuracy.

**Table 3:** RMISE over *g* ∈ [200, 30 000] (*g* ∈ [1 000, 30 000]) for five *N* (*g*) inference methods. RMISE = root mean integrated squared error (in units of 10^5^). M = 10^6^, K = 10^3^. Values are averages over the 25 curves of Fig. 2.

| Model | InferNo |  | Relate |  | PSMC |  | Mushi |  | CHIMP |  |
| --- | --- | --- | --- | --- | --- | --- | --- | --- | --- | --- |
| 1 | **1.8** | **(1.7)** | 7K | (9.9) | 722 | (4.3) | 682 | (2.0) | 7.7 | (7.6) |
| 2 | **2.0** | **(1.7)** | 14K | (8.6) | 0.9M | (10.1) | 1K | (8.6) | 0.2M | (0.2M) |
| 3 | **3.5** | **(3.5)** | 4K | (9.5) | 9.5 | (6.1) | 289 | (8.1) | 9.8 | (9.8) |
| 4 | **5.5** | **(5.3)** | 9K | (9.4) | 3M | (7.3) | 769 | (7.0) | 8.0 | (7.8) |
| 5 | **7.6** | **(7.2)** | 6K | (17.2) | 4K | (10.8) | 343 | (9.8) | 16.6 | (14.9) |
| 6 | **7.7** | **(7.6)** | 15K | (10.3) | 5M | (8.4) | 3K | (49.1) | 9.1 | (8.9) |

**Table 4:** RMISE (in units of 10^5^) for InferNo and PSMC when *n* = 2. Values are averages over the 25 curves of Fig. 9.

| Generations | Model: | 1 | 2 | 3 | 4 | 5 | 6 |
| --- | --- | --- | --- | --- | --- | --- | --- |
| [200, 30 000] | InferNo | **8** | **6** | **13** | **16** | **21** | **14** |
|  | PSMC | 6M | 9M | 3M | 9M | 8M | 6M |
| [1 000, 30 000] | InferNo | **7.3** | **6.1** | **11.7** | **15.2** | **18.2** | **12.2** |
|  | PSMC | 30K | 3.0M | 14.3 | 2.7M | 51.0 | 900 |
| [5 000, 30 000] | InferNo | **5.2** | **5.4** | **9.0** | 7.8 | **11.6** | 8.6 |
|  | PSMC | 5.9 | 6.0 | 12.0 | **4.4** | 11.8 | **7.4** |

**Table 5:** Growth rates (per 10^5^ generations) for eight simulation models. Although increasing *g* corresponds to backwards in time, growth is measured forward in time so a positive growth rate means that *N* (*g*) decreases as *g* increases. The rates in each column apply from the generation shown at the top of that column until that shown in the next column, or for all *g >* 30*K* in the case of the final column. See Table 1 for population sizes at the same time points.

| Model | Time before present in units of $10^3$ generations | | | | | | | | | | | | |
| --- | --- | --- | --- | --- | --- | --- | --- | --- | --- | --- | --- | --- | --- |
|  | 0 | 0.025 | 0.03 | 0.3 | 0.4 | 1 | 1.8 | 1.9 | 3 | 9 | 18 | 18.1 | 30 |
| 1 | 0 | 0 | 0 | 0 | 0 | 0 | 0 | 0 | 0 | 0 | 0 | 0 | 0 |
| 2 | 10 | 10 | 10 | 10 | 10 | 10 | 10 | 10 | 10 | 10 | 10 | 10 | 0 |
| 3 | -50 | -50 | -50 | -50 | -50 | -50 | -50 | -50 | -4 | -2 | 0 | 0 | 0 |
| 4 | 20 | 20 | 20 | 20 | 20 | 20 | 20 | 20 | 10 | -3 | -3 | -3 | 0 |
| 5 | 0 | 0 | 0 | 1 400 | 0 | 0 | -1 200 | 0 | 0 | 0 | 500 | 0 | 0 |
| 6 | 700 | 700 | 700 | 700 | -270 | 0 | 0 | 0 | 27 | -15 | 0 | 0 | 0 |
| 7 | 20 000 | 0 | 0 | 0 | 0 | 0 | 0 | 0 | 0 | 0 | 0 | 0 | 0 |
| 8 | 5 000 | -15 000 | 10 | 10 | 10 | 10 | 10 | 10 | 10 | 10 | 10 | 10 | 10 |

**Table 6:**
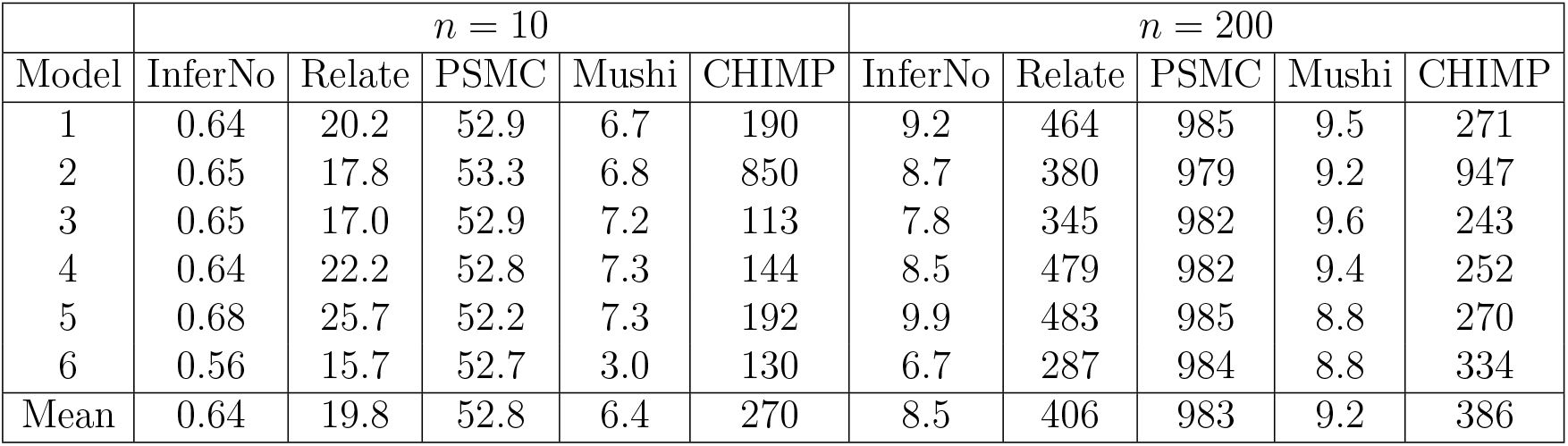
CPU time (seconds). Average computing time (over the 25 replicates in each setting) for the results in Figure 8 (*n* = 10) and Figure 3 (*n* = 200).

InferNo is also the fastest among the five methods (Table 6), taking on average (over 25 replicate datasets for each of 6 models) 8.5 seconds per analysis when *n* = 200, just faster than 9.2 seconds for Mushi and much faster than 386 seconds for CHIMP, 406 seconds for Relate and 983 seconds for PSMC. Mushi has a computing cost similar to InferNo when *n* = 200, and is slower when *n* = 10.

Increasing *n* improves inference more for recent generations (*g <* 1 000) than the distant past (Fig. 10), because many lineages coalesce in recent generations. The average RMISE for *g* ∈ [200, 30 000] decreases by 82% when *n* increases from 10 to 1 000 (Table 7), but when restricted to *g* ∈ [1 000, 30 000] RMISE decreases by only 39%. Table 7 and Fig. 11 show little effect on inference of misspecifying recombination rates, with average RMISE increasing 15% or 18% when the recombination map of Fig. 1 or a constant rate 50% higher than that assumed by InferNo is used for the data simulation. InferNo is also robust to sequencing errors, with RMISE increasing by 9%, 20% and 29% when sequencing errors are introduced at rate 1, 2 and 3 errors per 10^5^ sites (Table 7 and Fig. 12).

**Table 7:**
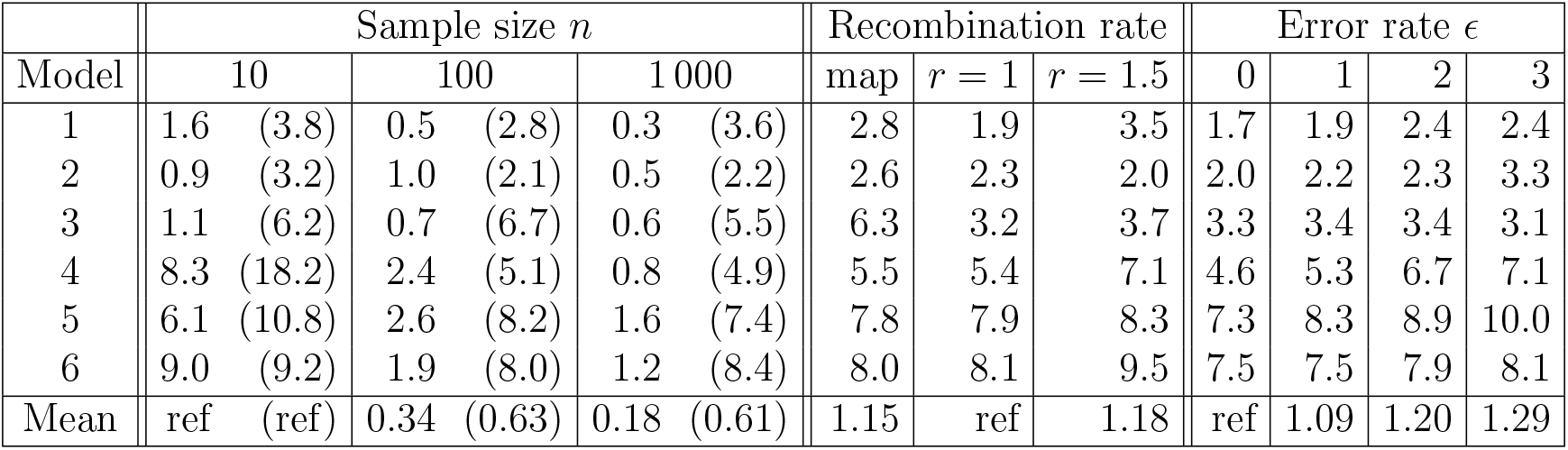
RMISE of InferNo (in units of 10^5^) under perturbations of the data generating model. Entries in the bottom row are expressed relative to a baseline indicated by “ref”. *r* is stated in units of 10^−8^ and *ϵ* is per 10^5^ sites. Entries under “Sample size” correspond to the curves in Figure 10, with integration over *g* ∈ [200, 1 000] (*g* ∈ [1 000, 30 000]). Entries under “Recombination rate” correspond to the curves in Figure 11 and those under “Error rate” correspond to the curves in Figure 12; in both cases the integration is over *g* ∈ [200, 30 000].

Because the computing time is approximately linear with *n* (Fig. 10), InferNo can exploit large sample sizes to obtain accurate estimates of *N* (*g*) in the recent past (small *g*). Fig. 3 (top) demonstrates its capability for accurate estimation of recent population sizes, in comparison with IBDNe (middle). The RMISE for IBDNe is about 20-fold higher than for InferNo (638 vs 32). With *n* = 2×10^5^ (Fig. 3, bottom), InferNo required only 326 seconds for each analysis to accurately infer rapid size changes over the past 30 generations. It is feasible to use InferNo on even the largest available human biobanks to infer changes in population size over recent centuries, which may be due to migration events, plagues or environment changes.

**Figure 3.**
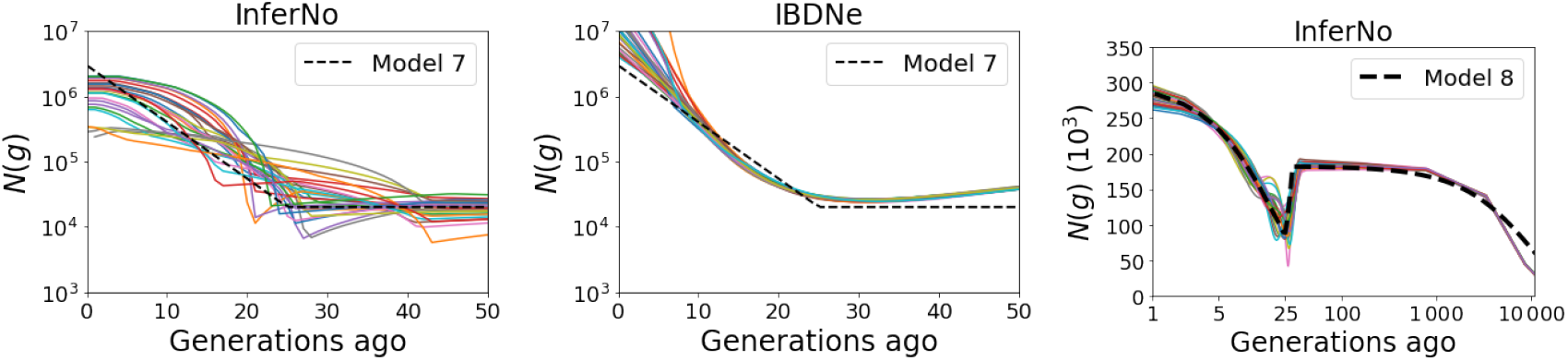
InferNo and IBDNe estimates of *N* (*g*) under Models 7 and 8. The sample size and sequence length for Model 7 (top and middle panels) are *n* = 2 000 and *ℓ* = 10^8^ sites. For Model 8 (bottom), they are *n* = 2×10^5^ and *ℓ* = 5×10^7^ sites.

### Results from human data analysis

When presenting results for human demographic history, for consistency with other authors we will express time as years (*t*) rather than generations (*g*) before present. We will also use K as shorthand for 10^3^. Unfortunately there is no agreement among authors on generation time; we adopt a recent estimate of 27 years per generation (Wang et al., 2023).

Since *N* counts haploid genomes, the number of humans is *N/*2.

For *t >* 25K there is a striking similarity of the inferred effective population size curves cross the four populations within each region (Fig. 4), despite the InferNo analysis being performed independently for each population. The four AFR populations have *N* (*t*) *>* 40K for *t <* 350K, while only for 550K *< t <* 600K are all four *N* (*t*) values below 20K. The lowest *N* (*t*) in any AFR population was 17.6K in MSL at *t* = 590K. The 12 non-AFR populations have *N* (*t*) *<* 40K for 30K *< t <* 500K.

**Figure 4.**
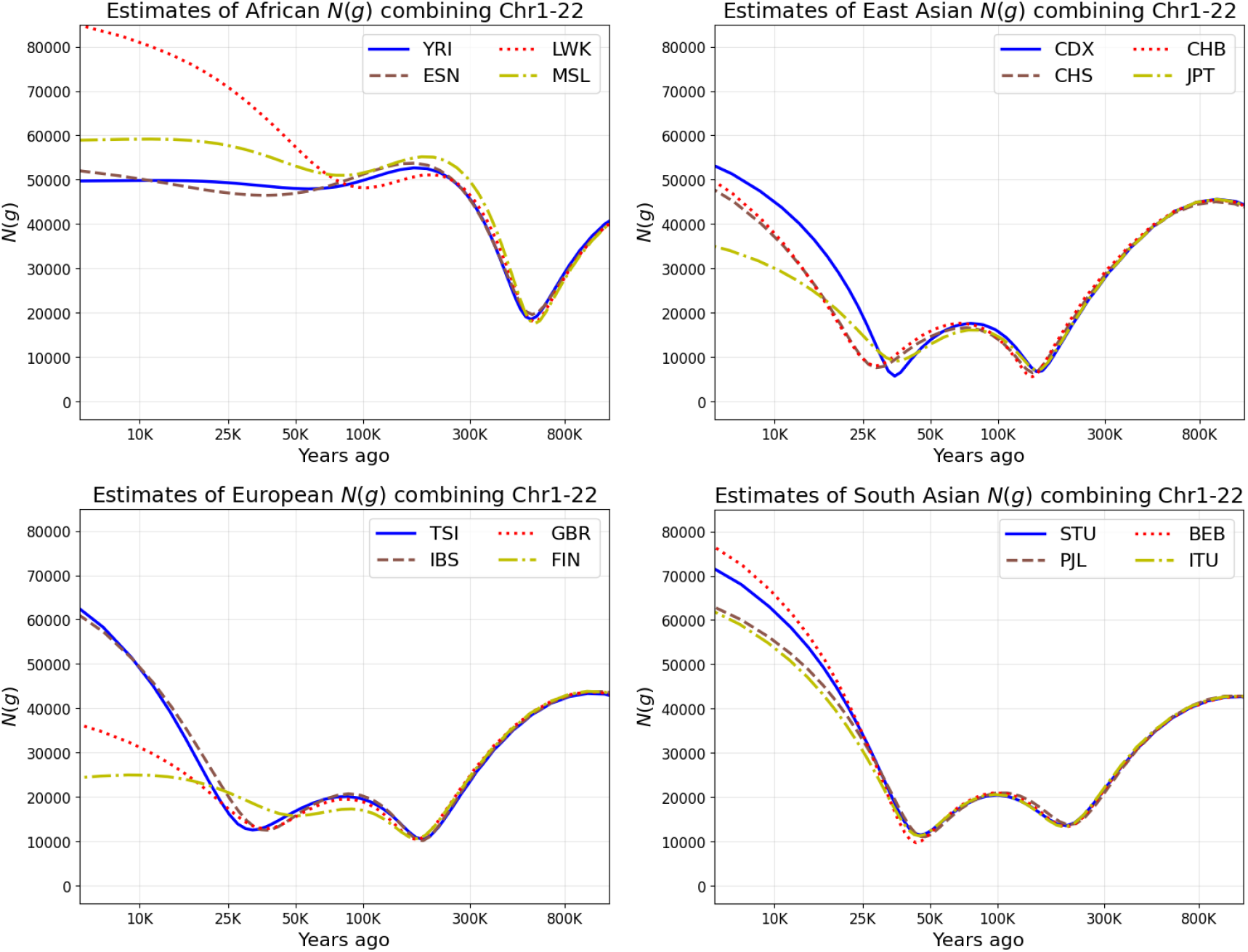
InferNo estimates of population size *N* for four populations in each of four regions. See Table 2 for the population codes. The time axes are on a logarithmic scale.

The population sizes of AFR and non-AFR populations remain distinct at *t* = 800K, estimated at 28K – 30K for all 4 AFR populations and at 41K – 45K for all 12 non-AFR populations. This time is well before the migration of modern humans out of Africa, which is typically estimated around *t* ≈ 50K Ragsdale et al. (2023). Estimates of current size *N* (0) are much more variable, ranging from 21K in FIN to 88K in LWK.

The non-AFR populations each experienced two local minima (bottlenecks). The most severe bottleneck arose in the ancestry of the EAS populations when *t* ≈ 135K, with *N* (*t*) estimates between 5.6K (CHB) and 7.1K (JPT). The corresponding bottleneck in EUR populations was inferred to be less severe (between 10.2K (IBS) and 10.7K (FIN)) and earlier (*t* ≈ 175K). For the SAS populations, the more recent of the two bottlenecks (*t* ≈ 45K) was slightly more severe than the earlier bottleneck (*t* ≈ 195K), ranging from 9.7K (BEB) to 11.4K (STU). The more recent bottleneck was even more recent in EAS (28K *< t <* 36K), with *N* (*t*) ranging from 5.7K (CDX) to 9.1K (JPT). In EUR, the more recent bottleneck was negligible in FIN but arose at a similar time to EAS in the other three populations (32K *< t <* 37K), with *N* (*t*) ranging from 12.5K (IBS) to 12.8K (GBR).

Fig. 13 shows, for 4 of the 16 populations, that the major features of demographic history inferred by InferNo are detected using each type of statistic individually: AFS only (*ω* = 0) and TMRCA estimates only (*ω* = ∞), but with some differences in timing and severity of bottlenecks, particularly for the older bottleneck in the two non-AFR populations. These discrepancies reflect uncertainty in inferences arising from modelling assumptions that inevitably deviate from the complex reality of human history.

As a further check on our inferences about human demographic history, we performed two further simulation experiments. First, we simulated a dataset with *n* = 200 and an *N* (*g*) curve similar to those inferred for the non-AFR populations. We then applied InferNo using the same *λ* value as for the real-data analysis. Fig. 5 (left) shows that InferNo accurately recovers the curve, which is also the case when the experiment is repeated for an AFR population (Fig. 14). We repeated the non-AFR simulation now including an additional, ancient bottleneck similar to that inferred for the AFR populations. Fig. 5 (right) shows that InferNo captures much of the signal from the ancient bottleneck, and that inference of the more recent bottlenecks remains accurate.

**Figure 5.**
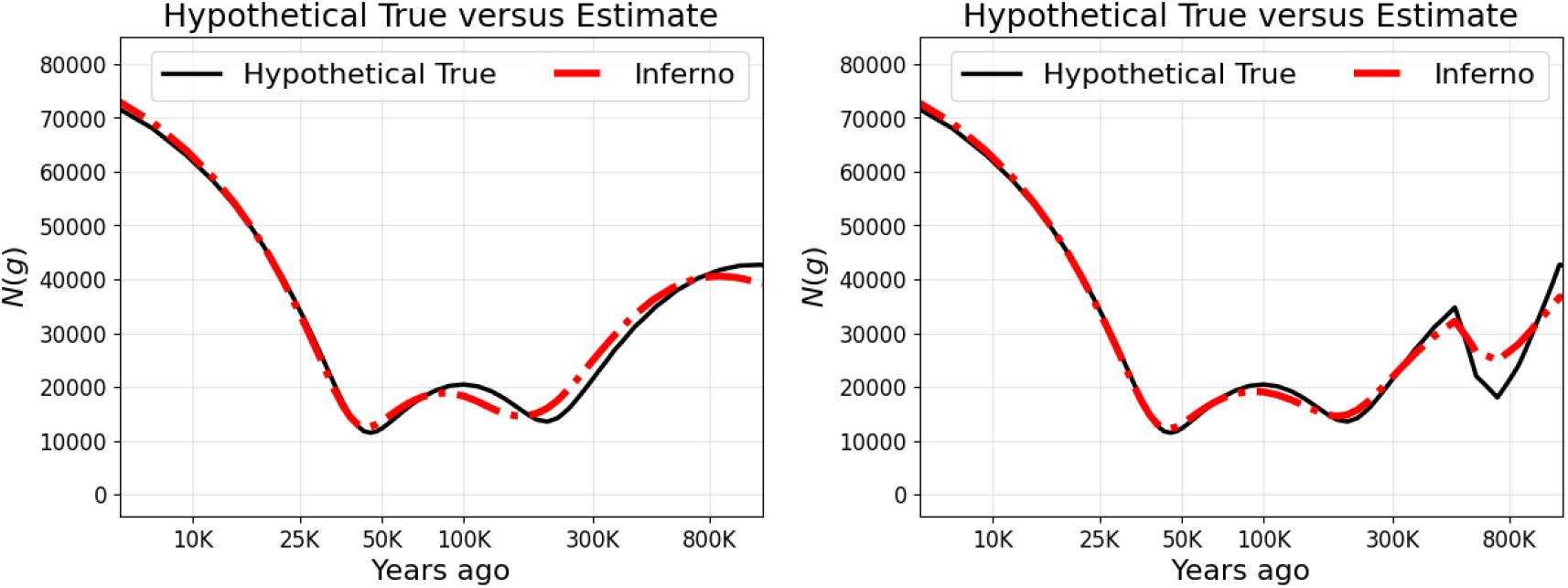
Testing InferNo inferences from human data. Left: data are simulated with *N* (*g*) (black curve) close to those inferred for the non-AFR populations (Fig, 4), and then InferNo used for inference (red curve). See Fig. 14 for the corresponding plot for AFR populations. Right: we add an ancient bottleneck similar to that inferred for the AFR populations, and are able to recover it in the inference. The time axes are logarithmic. Both simulations generated data for all 22 autosomes with *n* = 200, mutation rate *µ* = 1.3 × 10^−8^ and recombination rate *r* = 10^−8^.

## Discussion

InferNo achieves higher statistical and computational efficiency than existing approaches for estimating effective population sizes by combining analytical steps with simulation-based approximation. The constant-size simulation model is equivalent to the target variable-size model under a time scaling that is inferred from the observed AFS and (if required) from TMRCA estimates for sequence pairs in genome segments. This approach allows InferNo to achieve more precise population-size inferences, identifying features that are not detected by other methods.

We showed robustness of InferNo to sequencing errors and gene conversions, variation in the mutation rate along the genome, and misspecification of the recombination map. The better performance of InferNo over four alternative approaches is particularly marked for recent times (*g <* 1 000 generations), and it also outperforms an IBD-based method that focusses on the recent past. InferNo is the most computationally efficient of the methods, with cost increasing linearly with *n*. We illustrated its ability to accurately infer very recent population sizes from biobank-scale datasets.

We used InferNo to estimate historical population sizes for 16 worldwide human populations. The four populations within each of four worldwide regions showed a striking similarity of the inferred *N* (*t*) curves for *t >* 25K years before present. As expected, we found the greatest differences between African (AFR) and non-AFR populations, with these differences persisting more than 800K years into the past.

Our results show broad similarity with, but also important differences from, results of analyses of three of the populations studied here (Upadhya and Steinrücken, 2022, Fig 11), and different populations in similar regions (Bergström et al., 2020, Fig 4) using MSMC2 (Schiffels and Wang, 2020), which is related to PSMC and was found by Upadhya and Steinrücken (2022) to be inferior to their CHIMP software. Both these studies identified one long-duration bottleneck in the histories of EUR and EAS populations, with *N <* 20K for *t* between about 15K and 125K, with minima at *t* ≈ 38K Bergström et al. (2020) and *t* ≈ 100K Upadhya and Steinrücken (2022). There is no ground truth available to conclude that any method is most accurate, but we point to the better performance of InferNo in simulations, including under Model 6 with two bottlenecks. InferNo also infers a single bottleneck when its smoothing parameter *λ* is increased from the minimum-BIC value; however, the consistency in the timing of the two bottlenecks across populations in each region is evidence against them being statistical artifacts.

An ancient, severe bottleneck reported in ancestral AFR populations was not evident in analyses of non-AFR populations Hu et al. (2023). Our inferred AFR bottleneck is much less severe (*N* ≈ 17K versus *N* ≈ 1 280), of shorter duration and less ancient (550K *< t <* 600K versus 813K *< t <* 930K). The findings of Hu et al. (2023) have been challenged by Deng et al. (2024a), who suggest that the severe bottleneck may be an artifact of AFS-based inference. Although we also use the AFS, Fig. 13 shows that InferNo inferences restricted to pairwise TMRCA are similar to AFS-based inferences. Further criticism of Hu et al. (2023) has come from Cousins and Durvasula (2025) who found a better fit using Mushi, inferring a bottleneck that is comparable in severity with the one we infer, but its date agrees with that of Hu et al. (2023).

We agree with Hu et al. (2023), and previous authors cited therein, in finding markedly different demographic histories of AFR and non-AFR populations long before the migrations of modern humans out of Africa. This difference could be explained by persistent structure in the populations ancestral to modern humans, as has been reported Ragsdale et al. (2023); Cousins et al. (2025), with modern AFR and non-AFR populations having a different ancestry mix from ancient populations. Alternatively, Hu et al. (2023) propose that the ancient bottleneck is also present in the demographic histories of non-AFR populations but its signal has been erased by a more recent bottleneck corresponding to out-of-Africa migrations. This was tested and considered implausible by Cousins and Durvasula (2025), and we also show that such an erasure is implausible for the case of the bottlenecks that we infer: Fig. 5 illustrates that InferNo would have extracted a signal if the more ancient bottleneck we inferred for the AFR populations also existed in the histories of the non-AFR populations.

InferNo does not report uncertainty in estimates, but users can gain an understanding of its magnitude from simulations (such as column 1 of Fig. 6) and from comparisons across similar populations (such as Fig. 4). In common with other methods for inferring *N* (*g*), InferNo assumes a single, unstructured population, and hence it infers an effective population size that may differ from census size, for example in the presence of population structure. The method can be extended to two or more populations at additional computational cost. While InferNo avoids inferring recombination events, it requires enough recombination to generate replication along the genome. InferNo allows *µ*(*s*) to vary over sites *s*, but like other methods, it assumes that each *µ*(*s*) is constant over time. Subject to the above caveats, InferNo is applicable to many species, and can benefit inferences of other evolutionary parameters, such as species divergence times.

**Figure 6.**
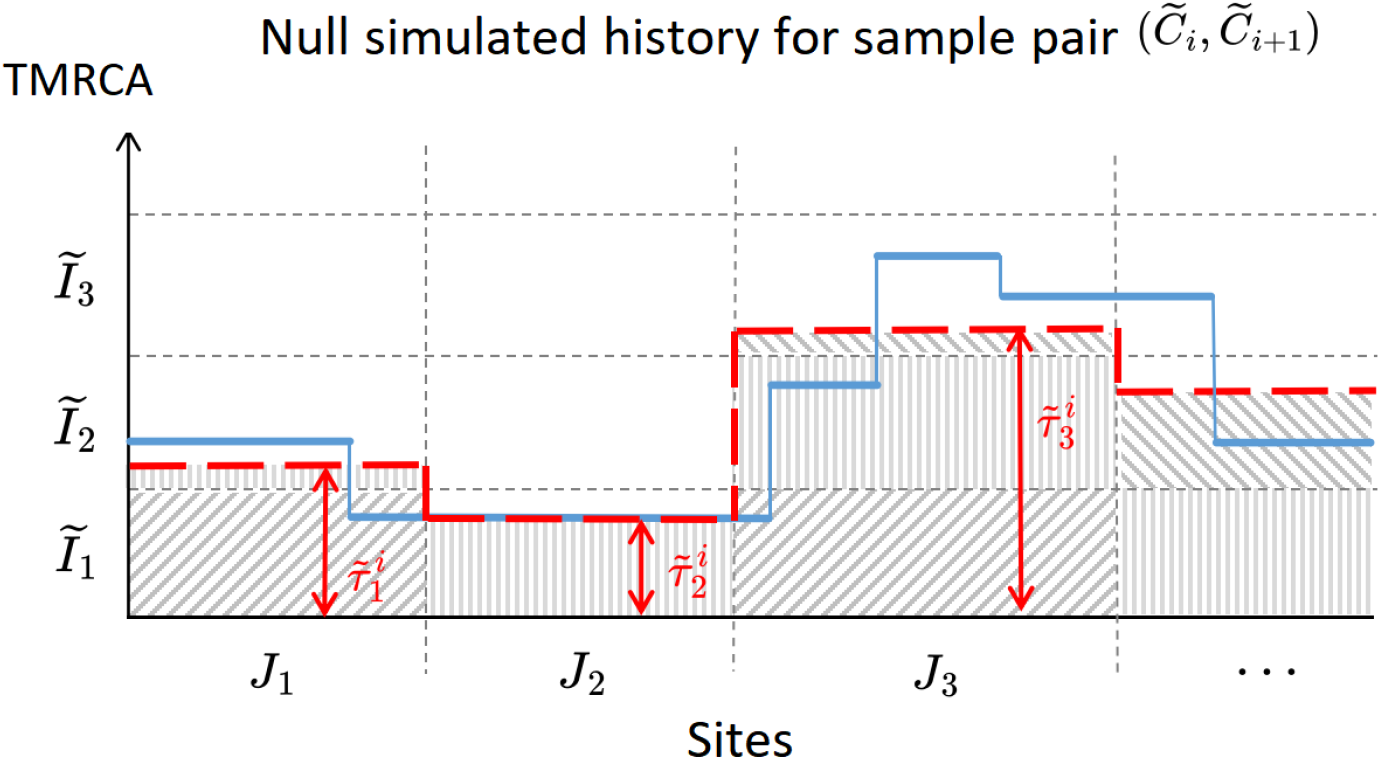
The TMRCA in a null simulation (solid blue) and its approximation 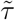 (dashed red) which equals the average blue value in each interval *J*_*l*_. The heights of the shaded rectangles correspond to the elements of 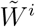 defined in the text.

The InferNo software and the data and code used in this paper are available at: github. com/ZhendongHuang/Inference_for_demographic_history.

## Supplementary Methods

### Inference of mutation rate

We estimate the mutation rate along the genome using the number of polymorphic sites close to any given site. More specifically, we partition the sites into *L*^∗^ segments 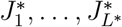, each of equal length *ℓ/L*^∗^, and estimate *µ*(*s*) by a piece-wise constant map fitted to 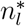, the number of polymorphic sites in 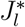:

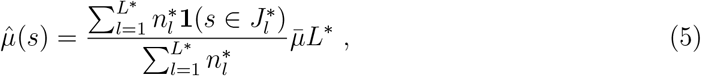

where **1**() is the indicator function.

### The null model and time scaling

Although there is flexibility in setting the null model population size, it should be not too far from average values over the historic periods of interest. We use 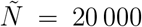. For large *ℓ* given the value of *r*, there is enough internal replication along the genome due to recombination that a single simulation is sufficient for precise estimation. We used one nullmodel simulation for all analyses reported here, but multiple simulations can be performed if desired for greater accuracy.

To relate the null model to the variable-size model, we consider the cumulative per-site coalescence rate over the interval *I*_*t*_ in the variable-size model, which is (*g*_*t*_ − *g*_*t*−1_)*/N* (*g*). Eq. [1] is obtained by equating this value to 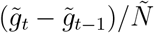, the corresponding quantity over 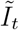 in the null model.

### Using the AFS

The {*k, t*} entry of 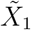 is the number of sites in the null simulation with a frequency-*k* derived allele for which the mutation arose during 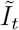. Because of the equivalence of null and size-varying models under the time scaling, the product of this entry with the time-scale ratio *β*_*t*_ equals in expectation the number of frequency-*k* sites in the size-varying model for which the mutation arose during *I*_*t*_. Summing over *t* gives the expected value of the *k*th entry of the AFS. Eq. [2] specifies this relationship jointly for each *k*.

### Using pairwise TMRCA estimates

From observed pairwise site differences, we estimate quantiles of the distribution of the TMRCA of pairs of observed sequences in short genome segments. We match these to the corresponding quantiles of the null model, obtained from simulated TMRCA values.

We choose the segments to obtain approximately *c* mutations per lineage per generation in each *J*_*l*_ by minimizing

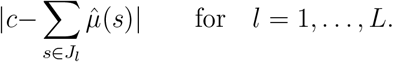

By default, we set *c* = 2.6 × 10^−4^, resulting in |*J*_*l*_| ≈ 20K sites for human populations, which is long enough to capture an informative number of site differences between sequence pairs, while limiting the within-segment variation in TMRCA. However, InferNo is robust to the choice of *c* because the same *J*_*l*_ are used for the null and variable-size models.

We write *τ* for the TMRCA of observed sequences *C*_*i*_ and *C*_*i*+1_ averaged over *s* ∈ *J*_*l*_ (see Fig. 6 for the corresponding averaging in the null model). To reduce computational effort, we only use neighbouring pairs of sequences (plus (*C*_1_, *C*_*n*_) when *n >* 2), so the resulting inferences can depend on sequence order; however, this effect is weak because every mutation is represented by a site difference in at least two neighbour pairs. By symmetry, *τ* has the same distribution for each *i*. We also treat the distribution as invariant over *l*, which holds approximately, and we make the corresponding assumption for the null model.

We use the sequence data to approximate the quantiles of the distribution of *τ*. Let *Z* denote an *n* × *L* matrix with {*i, l*} element equal to the number of site differences in *J*_*l*_ between the *i*th sequence pair. From coalescent theory we can model each element of *Z* as a Poisson(2*cτ*) random variable, conditional on *τ*. Let ℙ_*a*_(*b*) denote the probability mass function at *b* of the Poisson(*a*) distribution and *Z*_max_ = max{*Z*}. The probability mass function of each element *z* of *Z* is

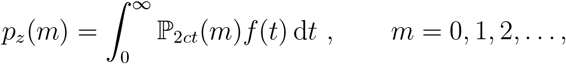

where *f* denotes the probability density function of *τ*. Discretising the integral at the points 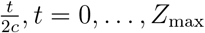, gives

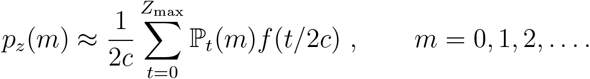

Restricting *p*_*z*_(*m*) to *m* ≤ *Z*_max_, defining *A* as a matrix with *A*_*i*+1,*j*+1_ = ℙ _*j*_(*i*), for *i, j* = 0, …, *Z*_max_, and defining a vector version of *f* evaluated at *t/*2*c*, for *t* = 0, …, *Z*_max_, gives the matrix equation

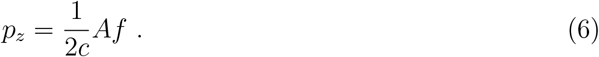

We estimate *p*_*z*_ by empirical values

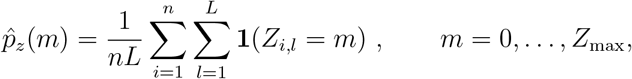

and then invert (6) to estimate *f* by

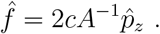

Let the vector 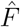 be the estimated cumulative distribution function of *τ* with *j*th element 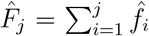 for *j* = 1, …, *Z*_max_+1. Then the *γ*-quantile of *τ*, for *γ* ∈ (0, 1), is estimated by linear interpolation: it is 0 if 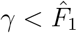, and

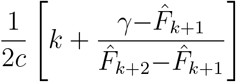

otherwise, where *k* ∈ {0, …, *Z*_max_−1} satisfies 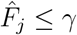 for *j* = 1, …, *k*+1 and 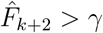. The *m*th element of the resulting length-*nL* vector *Y*_2_ estimates the *m/*(*nL*+1) quantile of *τ*.

We relate *Y*_2_ to the corresponding estimated quantiles for the null model, based on TMRCA values extracted from the null simulation. For a null-simulation sequence pair, let 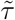 denote the average TMRCA over *J*_*l*_ (Figure 6). Let 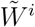 be an *L* × *T* matrix with {*l, t*} entry the contribution to 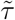 from the time interval 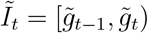,

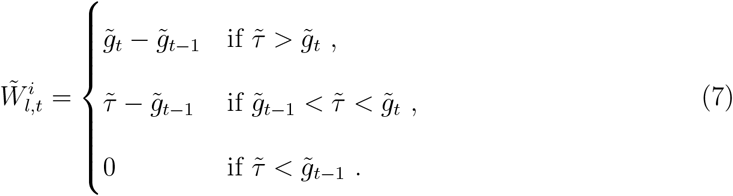

By construction, the sum of the *l*th row of 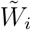 is the average TMRCA 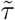.

Now let 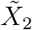 be the *nL*×*T* matrix constructed by vertically stacking the 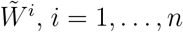, then sorting the rows to have increasing row sums. Since each row of 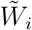 corresponds to a TMRCA, the rows of the sorted matrix correspond to the order statistics, which estimate the quantiles of 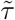. We can now infer *β* from regression equation Eq. [3], similar to that for *Y*_1_ and 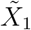.

### Combining the estimating equations

The first row of 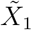 and of *Y*_1_ correspond to frequency-one alleles, which can be informative about recent *N* (*g*) but distorted by sequencing errors. By default, InferNo removes these first rows to improve robustness, but they can be retained if desired, for example when the sequencing quality is known to be high.

The matrix 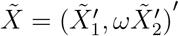 has dimensions *K*×*T*, where *K* = (*L*+1)*n*−2. For *n <* 2 000, we use the InferNo default 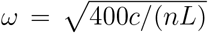, where *c* is the constant used to define the genome partition. Intuitively, this balances the amount of information from the AFS (represented by the sample size *n*) and the pairwise TMRCA (represented by *c/L*, approximately the expected number of mutations at each site). The constant 400 was found to work well for human data. Other choices for *ω* can be specified in InferNo, with higher *ω* tending to improve precision in the distant past at a cost to recent inferences. For *n* large we can set *ω* = 0, ignoring 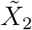 and *Y*_2_ which greatly reduces computational effort.

To infer *β* from the combined estimating equations (Eq. [4]), we weight the *k*th equation by an approximation to the standard deviation of *y*_*k*_, to give reduced weight to noisier observations. Let *S* be a diagonal matrix with *k*th diagonal element *s*_*k*_ = (*y*_*k*_+1)^−1*/*2^, *k* = 1, …, *K*, so that 1*/s*_*k*_ approximates the (Poisson) standard deviation of *y*_*k*_, adjusted to ensure finite values. We then estimate *β* by

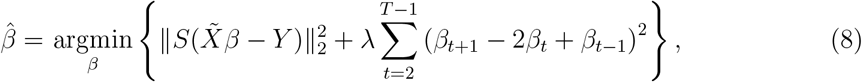

subject to *β*_*t*_ *>* 0, *t* = 1, …, *T*. The second term in (8) controls the smoothness of the *β*_*t*_ by penalising their second differences.

To select *λ*, we minimise the adjusted Bayesian information criterion (BIC) Schwarz (1978); Ye (1998):

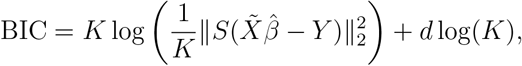

where *d* is the estimated degrees of freedom,

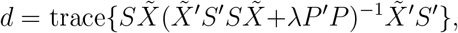

and *P* is a (*T*−2)×*T* matrix with entries *P*_*i,i*_ = *P*_*i,i*+2_ = 1 and *P*_*i,i*+1_ = −2 for *i* = 1, …, *T*−2, and *P*_*i,j*_ = 0 otherwise.

For simulation studies using Models 1 through 6, and for the real data analyses, we select *λ* to minimise BIC over log_10_(*λ*) ∈ {0,1,2,3,4,5,6,7}, although the extreme values 0 and 7 were never chosen in practice. We illustrate this approach for Models 4 and 5 with *n* = 100 (Figure 7). For Models 7 and 8, which focus on recent population size, BIC is computed using only the first ten rows of 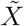 and *Y*, and the first ten rows and columns of *S*, which are informative about low-frequency alleles that predominantly arise from recent mutations. We select *λ* to minimise BIC over log_10_(*λ*) ∈ {−4, −3, −2, −1, 0} and log_10_(*λ*) ∈ {2.5,3,3.5,4,4.5} for Models 7 and 8, respectively.

**Figure 7.**
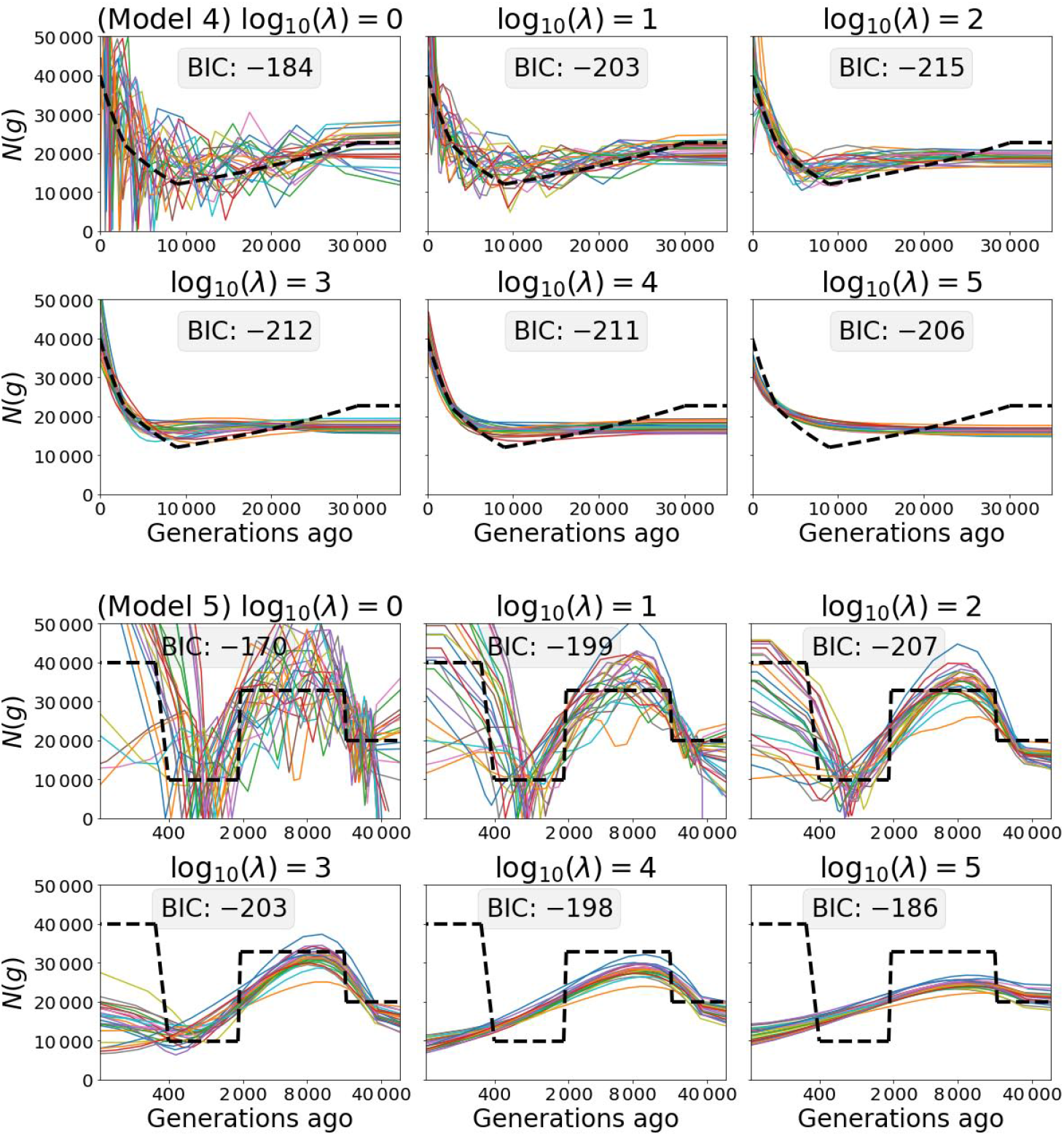
Selecting smoothing parameter *λ* using BIC (in units of 10^3^). Estimates of population size *N* (*g*), with *n* = 100, for Model 4 (top two rows) and Model 5 (bottom two rows, logarithmic time scale). The *λ* value is indicated above each panel. The minimum BIC is obtained for *λ* = 100 in both cases.

**Figure 8.**
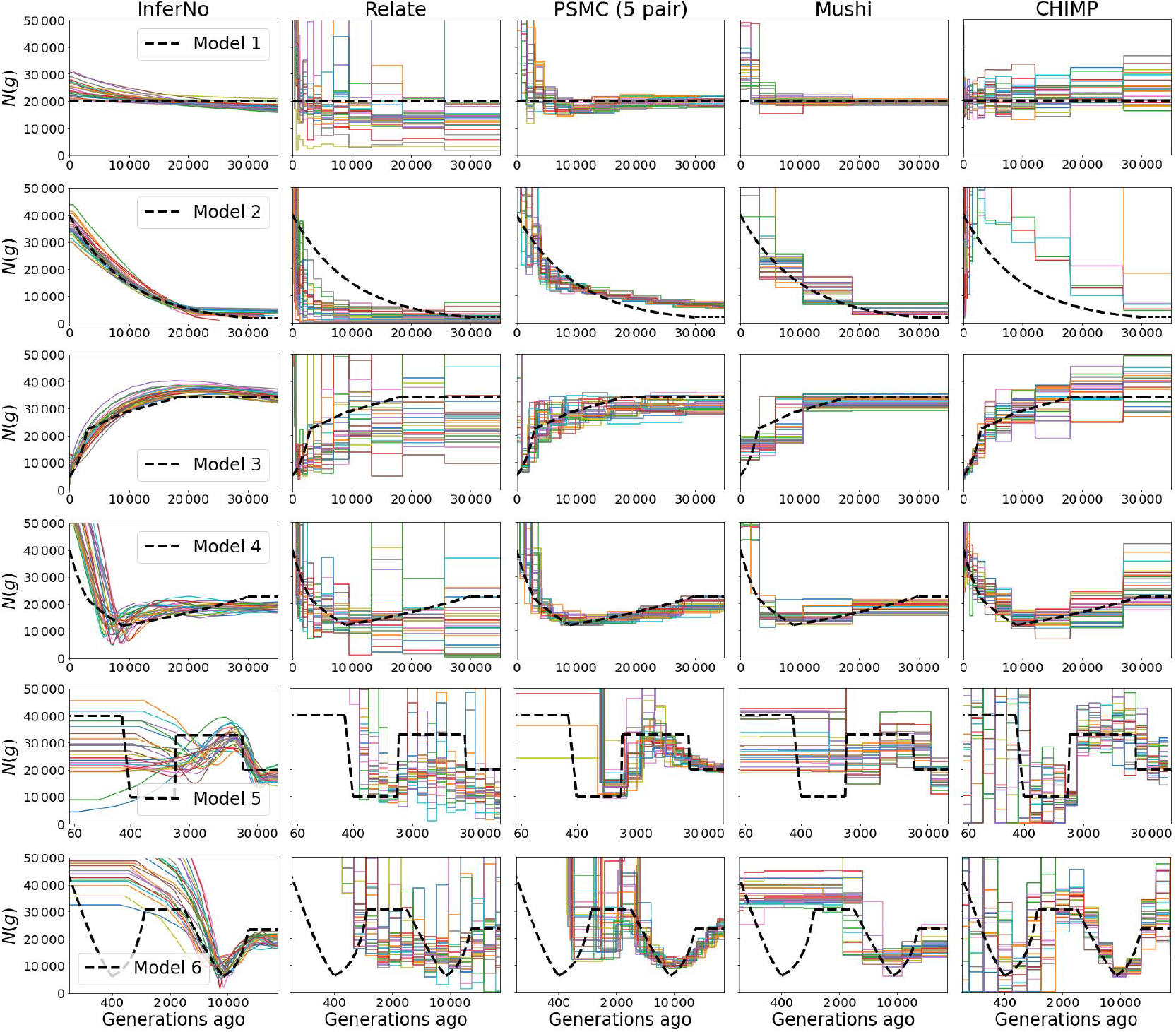
Comparison with other methods (*n* = 10). Other details are the same as for Figure 2.

**Figure 9.**
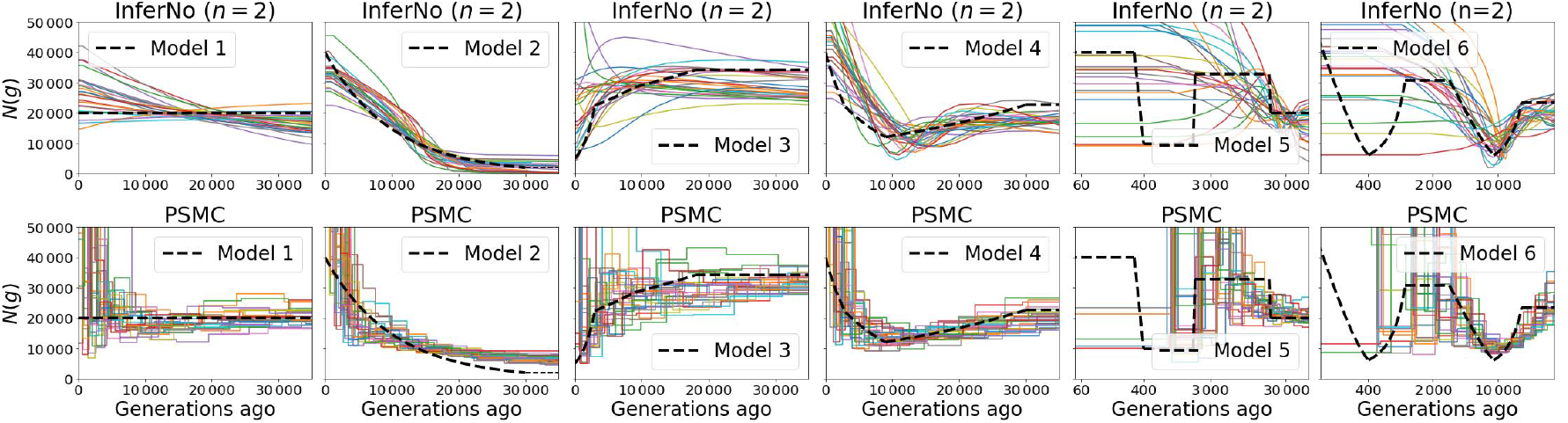
Comparison of InferNo with PSMC when *n* = 2. Sequence length *ℓ* = 10^7^ sites. Time scale is logarithmic for Models 5 and 6. See Table 4 for corresponding RMISE values.

**Figure 10.**
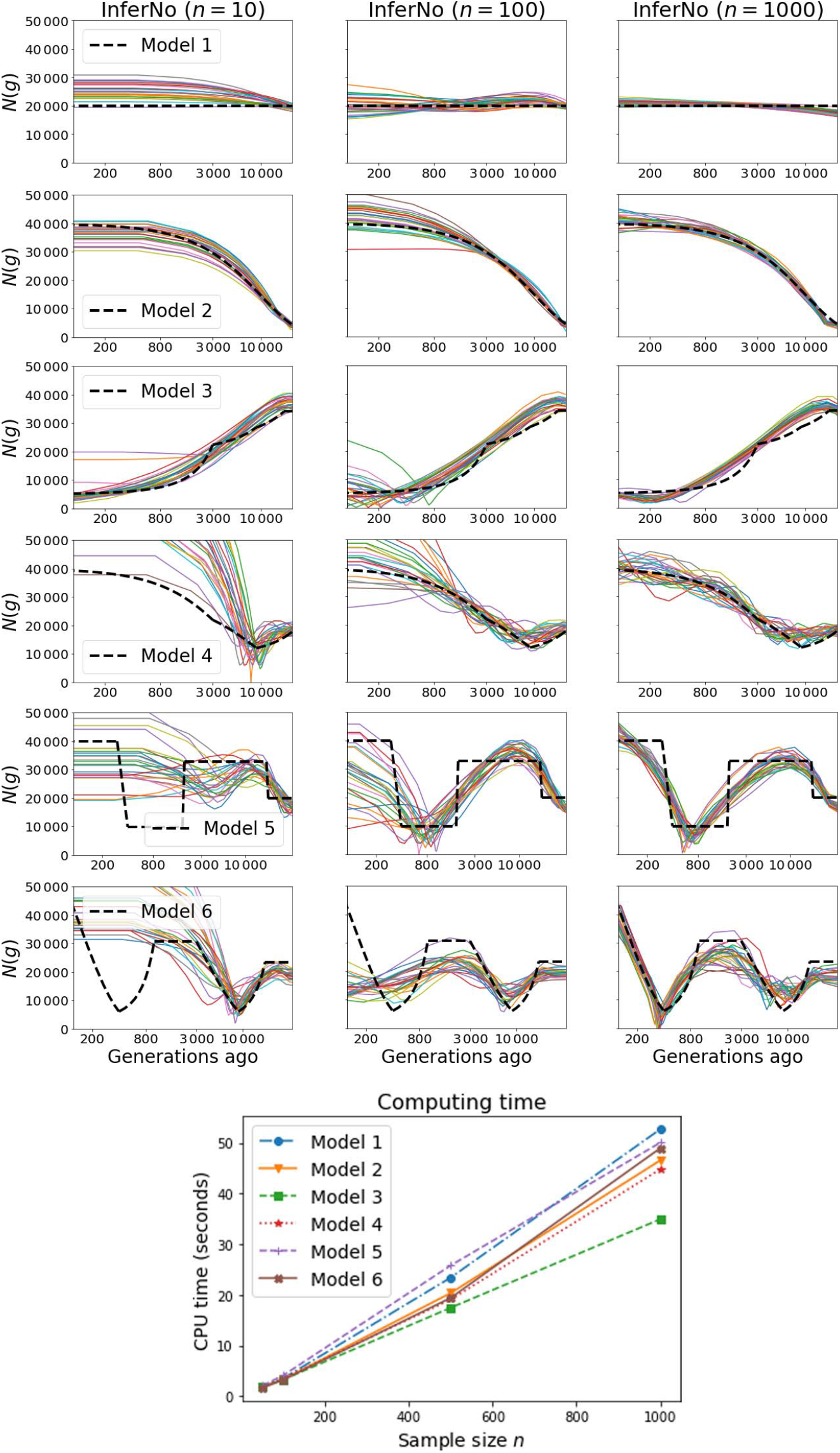
Estimation and computational performance with varying sample sizes. Rows 1-6: InferNo estimates of population size *N* (*g*) for three values of *n* in Models 1-6, with *g* on a logarithmic scale. Bottom: run time with *n* = 10, 50, 100, 500 and 1 000.

**Figure 11.**
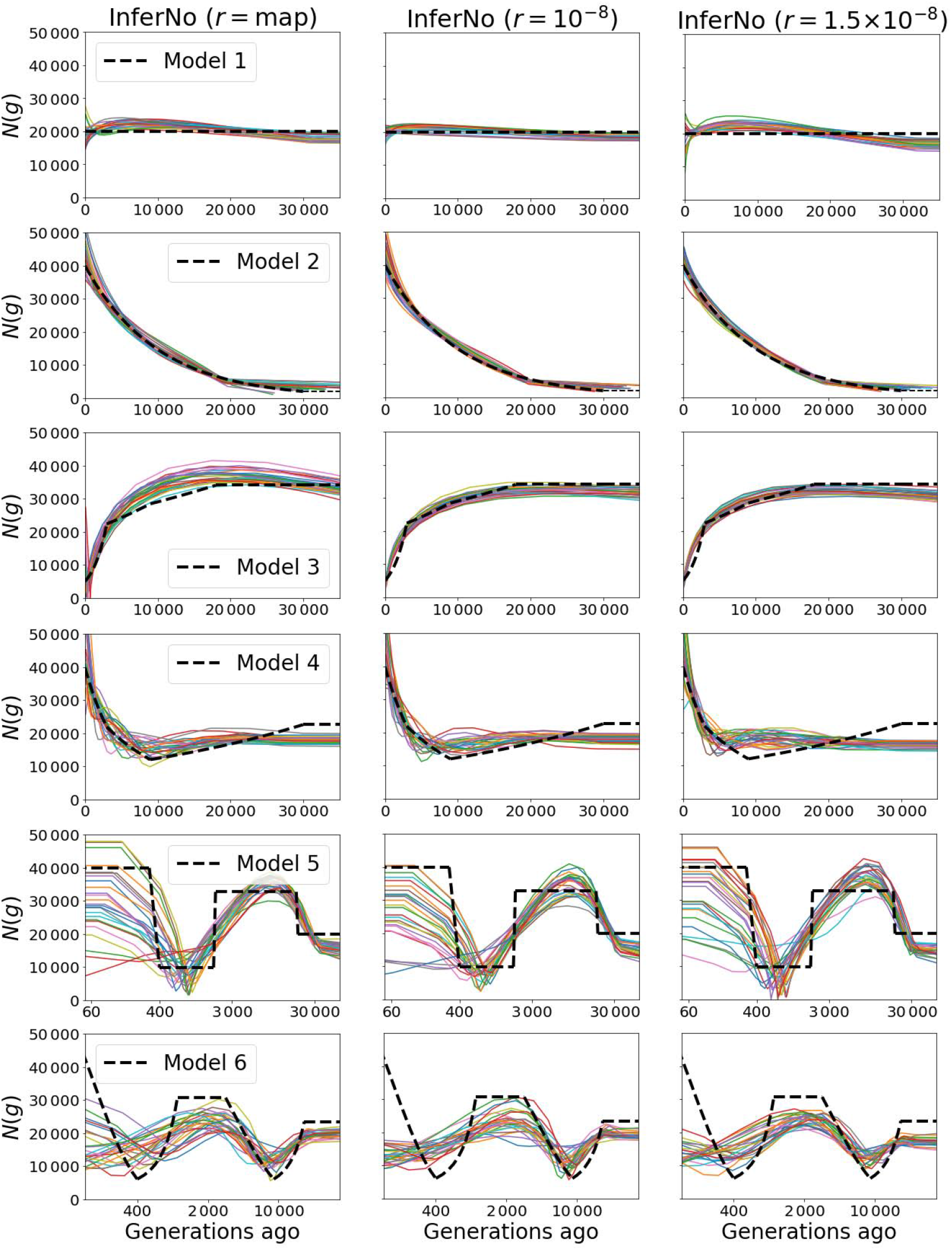
Performance of InferNo with misspecified recombination rates. Estimates of population size *N* (*g*) with *n* = 100 when the observed data is generated with the recombination map in Figure 2 (column 1), and with constant rates *r* = 1 × 10^−8^ (column 2) and *r* = 1.5 × 10^−8^ (column 3). The middle column corresponds to the recombination model assumed by InferNo. Rows correspond to simulation Models 1 through 6.

**Figure 12.**
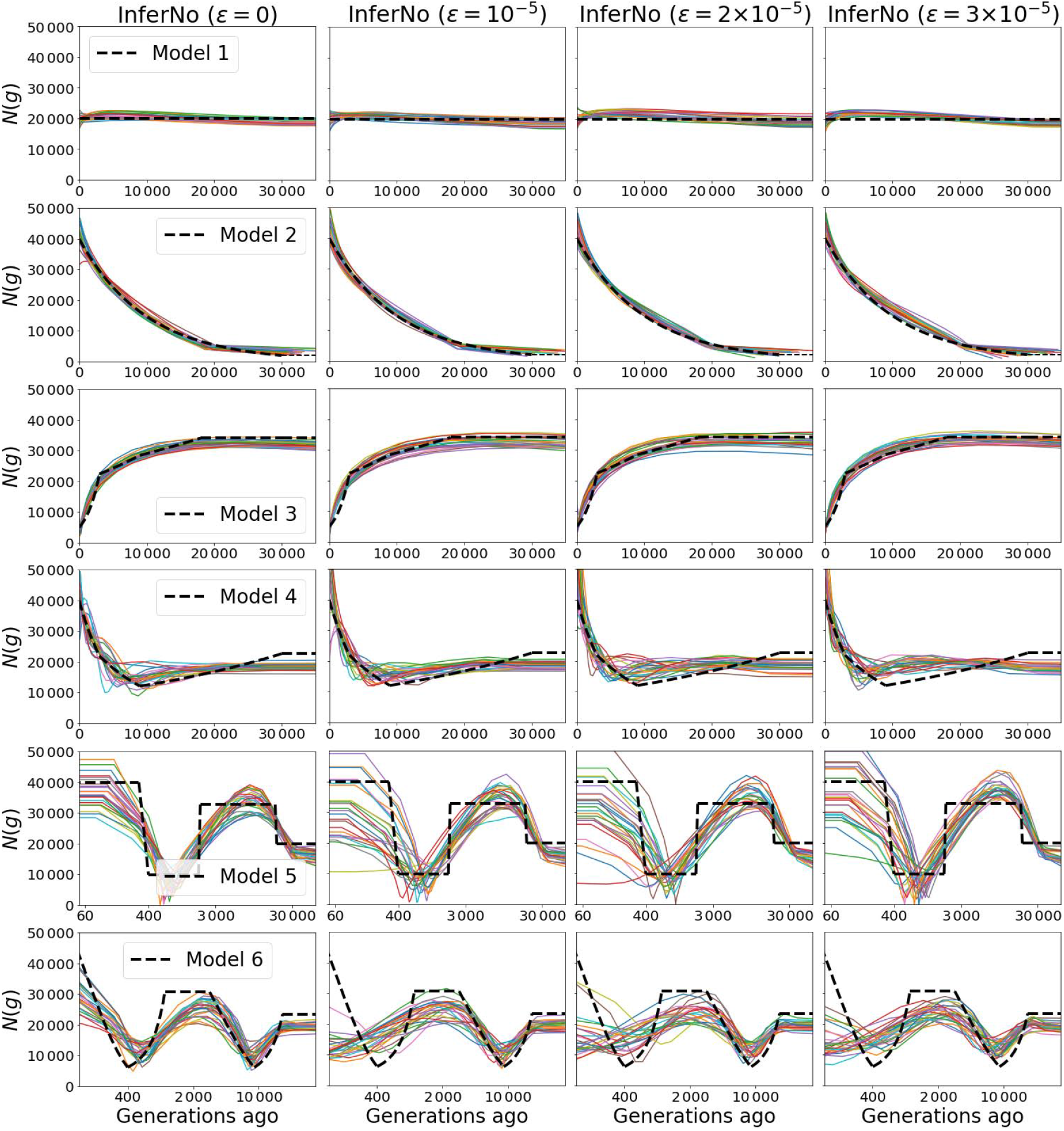
Performance of InferNo with different levels of sequencing error. Estimates of population size *N* (*g*) with *n* = 100. Columns correspond to sequencing error rates *ϵ* = 0, 1, 2, and 3 (per 10^5^ generations). Rows correspond to simulation Models 1 through 6.

**Figure 13.**
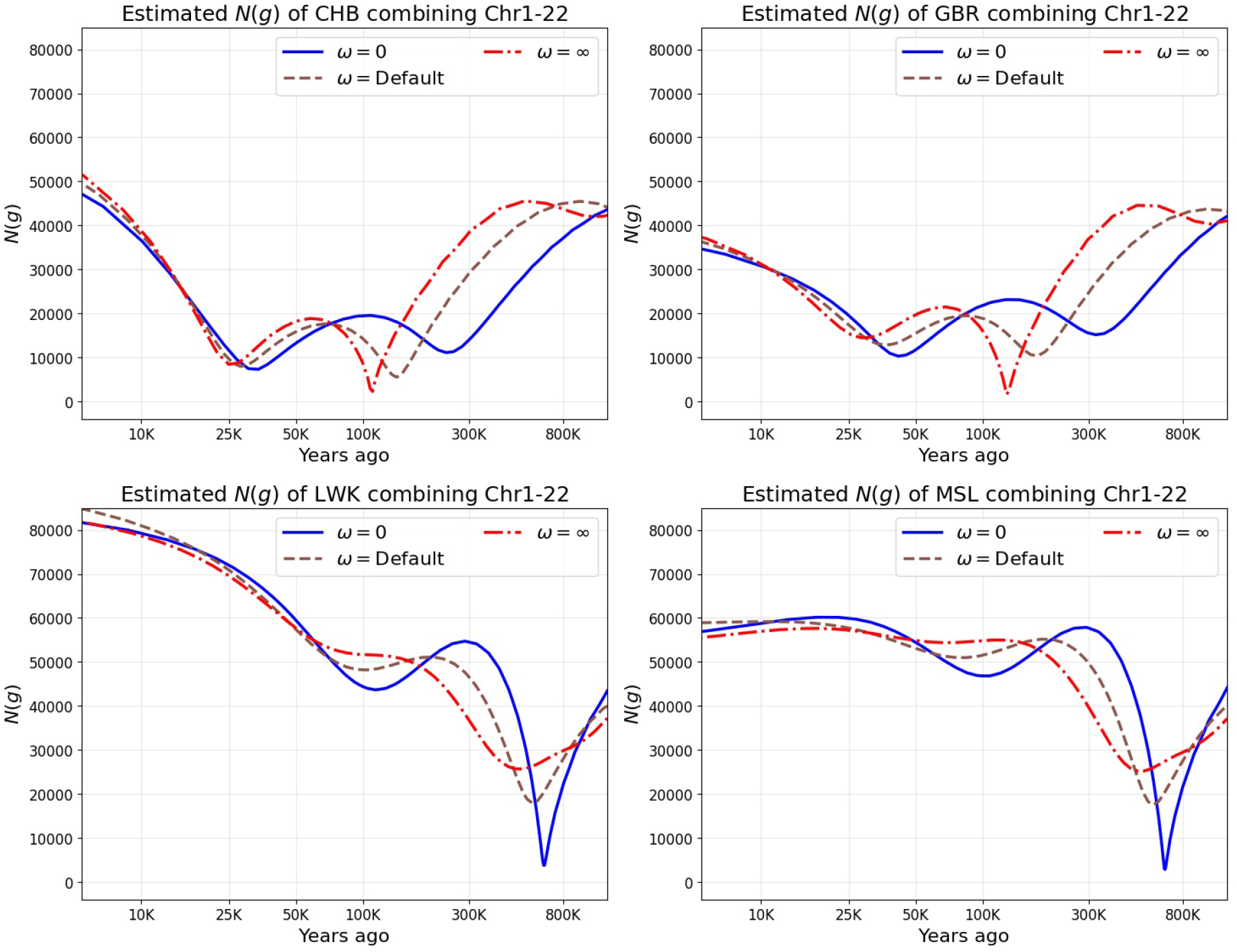
Components of InferNo inference for four human populations. Estimates of population size *N* (*g*) using only AFS (*ω* = 0), only TMRCA estimates (*ω* = ∞) and the default *ω* setting (used for all other plots).

**Figure 14.**
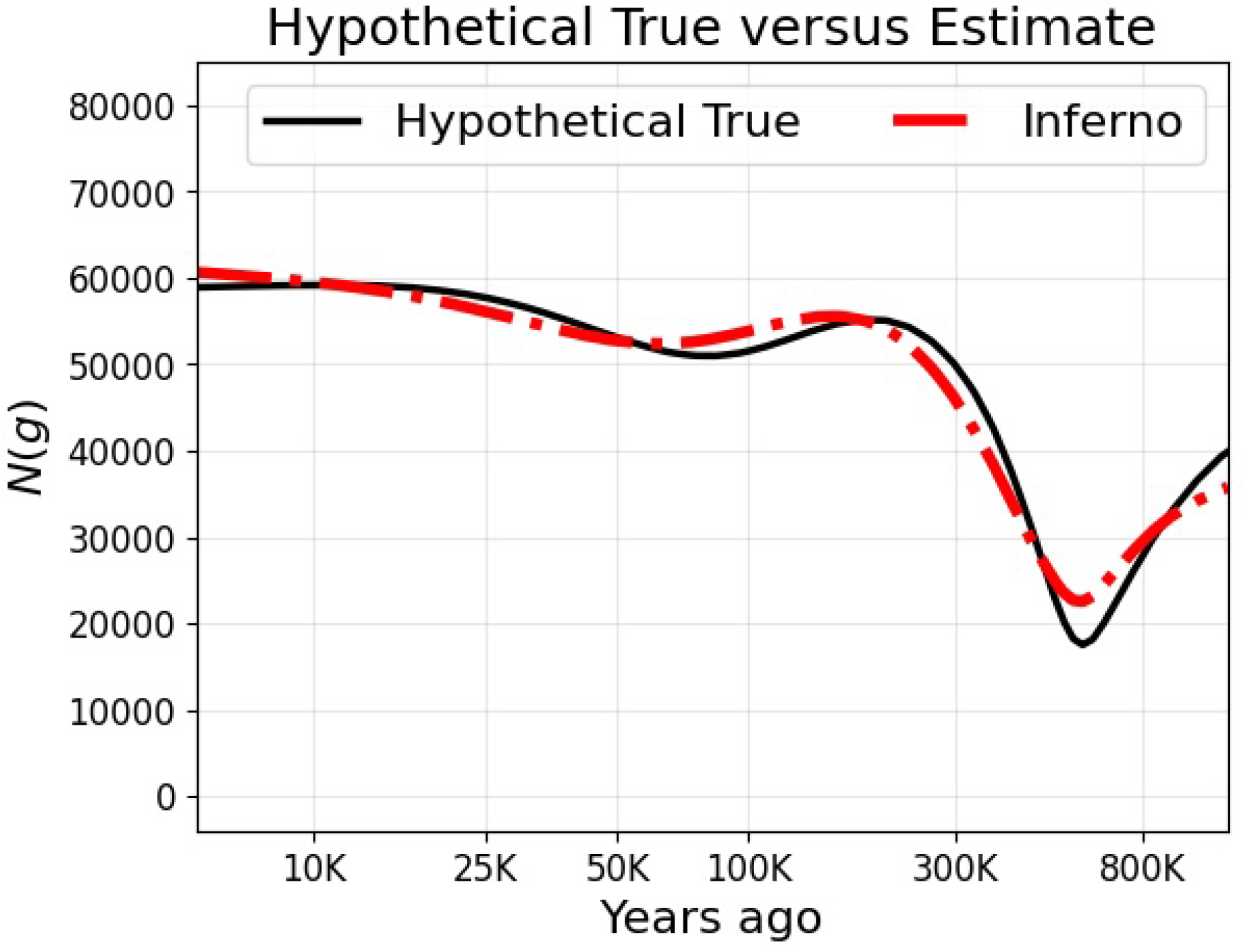
Testing InferNo inference for an AFR population. Similar to Fig. 5 (left) which shows inference for a non-AFR population. The red curve shows InferNo inference from *n* = 200 human sequences simulated with *N* (*g*) as shown in the black curve, similar to the inferred *N* (*g*) curve for AFR populations (Fig. 4, top left).

The final (optional) step is to convert the piece-wise constant estimate 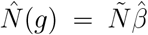 into a continuous, piece-wise-linear function with each interval midpoint remaining fixed, by connecting the points 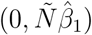 and 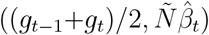 for *t* ≥ 1.

### Implementation details for other software

#### Relate

we set *N* (0) = 30 000 for both ARG inference and population size estimation.

#### PSMC

we set 20 free parameters, otherwise default settings were used. The original PSMC only uses a pair of sequences; for larger *n*, we implemented PSMC by partitioning the sequences into *n/*2 pairs and then averaging the resulting *n/*2 estimates.

#### Mushi

we set the maximum time investigated to 60 000 generations and the maximum number of iterations to 300. We obtain piece-wise constant estimates by fixing the trend order parameter to 0 and selecting the trend penalty parameter over {1, 3, 5, 10, 20, 40, 60, 100} in each setting. In practice, the extreme values 1 and 100 were never chosen.

#### CHIMP

we set `base` n = 2 and the right endpoints of the 20 time intervals as exp{3 + 7.6(*t*−1)*/*19} for *t* = 1, …, 20. To accelerate the convergence, we further initialise the population sizes at each time interval to be *N* (*g*) + *ε*, where *g* is the starting time of the interval and *ε* is a Gaussian error with mean zero and standard derivation *N* (*g*)*/*7. By doing so, we provided additional information of the demography, in return for faster computing time.

All other inputs are set by default. We use *T* = 20 time intervals for *N* (*g*) in all methods compared using Model 1-6, except for Relate, which automatically determines the time partition.

### Implementation details for human whole-genome data

Tree sequences data were downloaded from https://zenodo.org/records/5512994 and sequences extracted for both genomes of the 1 625 individuals whose population affiliation matched one of the 16 populations (Table 2).

We have previously shown Huang et al. (2025) that mutation rates can vary over chromosomes, and the pattern of variation may differ across populations. To allow for this variation, for each population we estimate the average mutation rate for chromosome *i* by 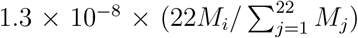 per site per generation, where *M*_*i*_ is the fraction of sites on Chromosome *i* that are polymorphic. Then we compute the matrices 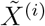 and *Y* ^(*i*)^ for chromosome *i*, and stacking them vertically to form 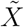 and *Y* for the InferNo inference.

